# Cell type- and layer-specific plasticity of olfactory bulb interneurons following olfactory sensory neuron ablation

**DOI:** 10.1101/2024.05.08.593238

**Authors:** Tenzin Kunkhyen, Taryn R. Brechbill, Sarah P.R. Berg, Pranitha Pothuri, Alex N Rangel, Ashna Gupta, Claire E.J. Cheetham

## Abstract

Lifelong neurogenesis endows the mouse olfactory system with a capacity for regeneration that is unique in the mammalian nervous system. Throughout life, olfactory sensory neurons (OSNs) are generated from olfactory epithelium (OE) stem cells in the nose, while the subventricular zone generates neuroblasts that migrate to the olfactory bulb (OB) and differentiate into multiple populations of inhibitory interneurons. Methimazole (MMZ) selectively ablates OSNs, but OE neurogenesis enables OSN repopulation and gradual recovery of OSN input to the OB within six weeks. However, it is not known how OB interneurons are affected by this loss and subsequent regeneration of OSN input following MMZ treatment. We found that dopaminergic neuron density was significantly reduced 7-14 days post-MMZ but recovered substantially at 35 days. The density of parvalbumin-expressing interneurons was unaffected by MMZ; however, their soma size was significantly reduced at 7-14 days post-MMZ, recovering by 35 days. Surprisingly, we found a transient increase in the density of calretinin-expressing neurons in the glomerular and external plexiform layers, but not the granule cell layer, 7 days post-MMZ. This could not be accounted for by increased neurogenesis but may result from increased calretinin expression. At subsequent time points, calretinin neurons in all three layers showed reduced density at 14 days but recovered to baseline by 35 days. Together, our data demonstrate cell type- and layer-specific changes in OB interneuron density and morphology after MMZ treatment, providing new insight into the range of plasticity mechanisms employed by OB circuits during loss and regeneration of sensory input.

## Introduction

Postnatal neurogenesis is rare in the mammalian nervous system. However, in the mouse olfactory system, multiple populations of neurons are generated throughout life from stem cells in two distinct locations: the olfactory epithelium (OE) and the subventricular zone (SVZ). This endows the olfactory system with substantial regenerative capacity and provides a unique opportunity to understand the mechanisms by which endogenously generated stem cell-derived neurons integrate into healthy and regenerating circuits in the brain.

The SVZ generates neuroblasts that migrate via the rostral migratory stream (RMS) to the olfactory bulb (OB), where they differentiate into multiple molecularly distinct sub-populations of interneurons located primarily in the glomerular layer (GL) and granule cell layer (GCL)^1–3^. Surprisingly, with the notable exception of the dopaminergic (DA) neurons, how most OB interneuron types are impacted by changes in sensory input from the OE is poorly understood. An early study found that naris occlusion in either neonatal or young adult rats reduced the density of calbindin-expressing (CB+) but not calretinin-expressing (CR+) periglomerular cells (PGCs) and reduced the density of parvalbumin-expressing (PV+) interneurons, which are found predominantly in the external plexiform layer (EPL)^4^. A more recent study found that naris occlusion in adult mice did not affect the survival of pre-existing CB+ or CR+ PGCs or the fate specification of newborn CB+ and CR+ PGCs^5^. Hence, there is somewhat conflicted evidence for activity-dependent changes, especially in PGCs.

In contrast, the effects of blocking sensory input on DA neurons have been studied extensively. DA neurons are found mainly in the GL but also scattered in the EPL and play multiple important roles in odor processing, including inhibition, gain control and decorrelation of odor representation in OB output neurons^6,7^, inhibition of periglomerular cells (PGCs) and dual inhibitory-excitatory modulation of external tufted cells surrounding distant glomeruli^8,9^, and reducing OSN release probability^10–13^. It has long been known that both dopamine production and expression of tyrosine hydroxylase (TH), the rate-limiting enzyme for dopamine biosynthesis, are strongly reduced when sensory input is blocked via either olfactory epithelium lesion or naris occlusion^14–17^. TH expression levels do recover to baseline levels after restoration of sensory input^15,18^. More recently, using Cre recombination-based labeling strategies that indelibly mark DA neurons and hence circumvent activity-dependent changes in TH expression, it has been shown that DA neuron survival is also reduced by a four-week naris occlusion^18,19^. However, the magnitude of DA neuron cell death varied widely between these two studies: 40 % in Sawada et al.^19^, vs. 5 % in Angelova et al.^18^. Nonetheless, both studies found that four weeks of restored sensory input was sufficient for DA neuron density to recover to baseline^18,19^, suggesting the incorporation of newly generated DA neurons. Notably, however, the effect of OE damage on DA neuron survival and neurogenesis remains unknown.

Sensory input to the OB is provided by olfactory sensory neurons (OSNs), which are themselves generated throughout life from basal stem cells located in the OE. OSNs can be generated from both globose and horizontal basal cells, with HBCs playing an important role following OE damage^20–22^. Notably, naris occlusion in adult mice does not alter OSN density or neurogenesis^23^, suggesting that the impact of naris occlusion vs. OE damage on OB interneurons may be very different. Methimazole (MMZ) treatment has emerged as an interesting model in which sensory input to the OB is abruptly lost and then gradually regenerates. A single dose of methimazole pharmacologically ablates OSNs while leaving basal stem cells unaffected, such that the olfactory epithelium is repopulated with OSNs within 4-6 weeks^24–26^. Regeneration of sensory input to the OB begins within a week, with the first immature OSNs generated after MMZ treatment providing sufficient sensory input to permit recovery of odor detection and even some simple odor discrimination behavior^27^. Over longer time periods, it has been shown that OSN inputs to external tufted cells return in 16 days^25^, while OSN projections to certain glomeruli and behavior in a previously learnt odor discrimination task recovers within 45 days^26^. Determining how MMZ impacts the size of OB interneuron populations can therefore provide important insight into their survival and the extent to which adult neurogenesis can restore depleted neuron populations.

Our goal in this study was to determine the effects of MMZ treatment on OB interneuron survival and regeneration, focusing on three interneuron subtypes. DA neurons were selected due to their strong dependence on sensory input for TH expression and survival and their ability to repopulate following naris occlusion. PV+ neurons were of interest because a large activity-dependent reduction in their density has been reported^4^, yet they are not normally generated in adults. Indeed, in mice, most PV+ neurons are generated during late embryonic and early postnatal life, with neurogenesis ceasing as early as postnatal day (P)7^28,29^, suggesting that lost PV+ neurons cannot be replaced. PV+ neurons form dense reciprocal connections with mitral and tufted cell dendrites^30–34^ and are thought to modulate OB output gain^32^; hence, changes in their density could have an important impact on OB output. Finally, CR+ neurons were selected because they are the most abundant subtype of adult-born OB neurons^28,35^, comprise both PGC and granule cell (GC) populations, and are generated not only in the SVZ but also in the core of the OB itself and possibly the rostral migratory stream ^36–41^.

Calretinin-expressing (CR+) periglomerular cells (PGCs) are the most numerous subtype of glomerular layer (GL) interneurons^42–44^ and are mostly generated postnatally^28,29,35,45^. They express an unusually restricted range of voltage-gated channels, resulting in very high input resistance and firing of at most one action potential in response to depolarizing current injection^46,47^. They also receive few synaptic inputs and may not form functional output synapses^47,48^. As a result, their function remains enigmatic, although it has been suggested that they improve signal:noise ratio at an early stage of olfactory processing by inhibiting principal neuron responses to asynchronous signals^46^. A significant fraction of superficial GCs also express CR; their neurogenesis peaks around birth but persists into adulthood^28,29^. CR+ GCs are morphologically similar to surrounding GCs, and receive similar excitatory inputs, but fewer inhibitory inputs than other GC types^49^. CR+ GCs are preferentially activated during odor discrimination and contribute to fine odor discrimination learning^49^.

We found that TH+ neuron density underwent bidirectional changes in the 5 weeks following MMZ treatment, validating the loss and subsequent regeneration of OB sensory input. While PV+ neuron density was unaffected by MMZ treatment, PV+ soma area was strongly reduced 7-14 days post MMZ but recovered within 35 days. Surprisingly, there was a transient increase in CR+ neuron density at 7 days post-MMZ only in the GL and EPL, while CR+ neuron density in these layers and the GCL was reduced by 14 days but recovered by 35 days post-MMZ. Together, our data demonstrate cell type- and laminar-specific changes in response to MMZ treatment, indicating the broad range of plasticity mechanisms, including but not limited to neurogenesis, available in the adult mouse OB.

## Methods

### Experimental animals

All procedures followed NIH guidelines and were approved by the University of Pittsburgh Institutional Animal Care and Use Committee. C57BL/6J mice (strain #000664) were purchased from the Jackson Laboratory and/or bred in-house, with new breeders purchased every 12 months to minimize genetic drift. Mice were maintained in individually ventilated cages at 22 °C and 48% humidity on a 12 h light/dark cycle with unrestricted access to food and water. Mice were group-housed unless same sex littermates were unavailable. A total of 71 mice were used in this study.

### Methimazole and EdU treatment

For analysis of neuronal density, 8-week-old C57BL/6J mice received an intraperitoneal injection of 75 mg/kg methimazole (MMZ (EMD Millipore)), dissolved in sterile saline, or an equivalent volume of sterile saline. Mice were perfused 7, 14 or 35 days after injection with MMZ or saline. For analysis of CR+EdU+ neuron density, 8-week-old C57BL6/J mice received an intraperitoneal (IP) injection of MMZ, then an IP injection of 50 mg/kg 5-ethynyl-2’-deoxyuridine (EdU (ThermoFisher Scientific)) 24 h later and were perfused after a further six days.

### Transcardial perfusion and cryosectioning

Mice were deeply anesthetized with 5% isoflurane in 1 l/min oxygen or with 200 mg/kg ketamine and 20 mg/kg xylazine, then transcardially perfused with ice-cold phosphate-buffered saline (PBS) and then 4% paraformaldehyde (PFA) in PBS. OBs were dissected and post-fixed overnight, then cryopreserved in 30% sucrose, embedded in 10% gelatin, and fixed/cryopreserved overnight in 15% sucrose/ 2% PFA. Gelatin blocks were flash-frozen in 2-methylbutane on dry ice and stored in a -80 °C freezer. 40 μm coronal sections were cut using a cryostat.

### Immunohistochemistry and EdU staining

One set of OB sections was used for staining with each primary antibody (to detect tyrosine hydroxylase (TH), calretinin (CR) and parvalbumin (PV), respectively). Free-floating sections were treated with 1 % sodium borohydride, washed with PBS, then blocked/permeabilized (5 % normal donkey serum [NDS]/0.5 % Triton X-100 in PBS) for 1h. Sections were incubated with primary antibody diluted in 3 % NDS, 0.2 % Triton X-100, 0.01 % sodium azide in PBS as shown in Table 1. After washing in PBS, sections were incubated in secondary antibody solution (Table 1; diluted in 3 % NDS, 0.2 % Triton X-100 in PBS) for 1 hour at room temperature. Sections were washed in PBS and mounted with Vectashield containing DAPI (Vector Labs).

OB sections from mice that had been injected with MMZ or saline and EdU underwent CR staining as described above, followed by EdU staining modified from Goedhart (2024)^50^. Briefly, these sections were permeabilized in 0.5% Triton X-100 in PBS for 30 min, then incubated with 200 mM copper sulfate, 4 mM sulfo-Cyanine5-azide (#A3330, Lumiprobe) and 114 mM ascorbic acid in PBS for 30 min in the dark. Sections were washed in 0.5% Triton X-100 in PBS and mounted.

### Widefield fluorescence microscopy

Tiled images of stained whole OB sections were acquired using either an Eclipse 90i microscope with a Plan Apo 10x/0.45 NA air objective, motorized stage and Elements Software (Nikon), or a Revolve microscope with Echo software (Echo), a 10x Plan Apo 0.4 NA air objective (Olympus) and Affinity Photo software (Affinity) for image stitching. Bandpass excitation/ emission filter wavelengths were (in nm): DAPI (385/30, 450/50); unstained green (470/40, 525/50); TH-, PV- or CR-AF546 (530/40, 590/50); EdU-AF647 (640/30, 690/50). We collected DAPI, green (unstained) and AF546 channels for analysis of TH+, PV+ and CR+ neuron density. We collected DAPI, CR-AF488, red (unstained) and EdU-AF647 channels for analysis of CR+EdU+ neuron density and CR fluorescence intensity. Detection settings were identical for all images collected for this data set.

### Image analysis

Image analysis was performed using Fiji (ImageJ)^51^. Anterior, central and posterior OB sections, which were approximately 25, 50 and 75 % along the anterior-posterior axis, were selected for analysis of TH+, PV+ and CR+ neuron density. OB layers were outlined in the DAPI channel of each section and used to demarcate regions of interest for cell counting or fluorescence intensity analysis. The area of each layer was measured, and neurons were counted manually within each layer using the multi-point tool. Neurons were not counted if they partially crossed the border with an adjacent layer. We confined our analysis to layers that had been shown previously to contain a significant density of TH+ (GL and EPL), PV+ (EPL, MCL/IPL, GCL) or CR+ (GL, EPL, GCL) neurons. We removed potential contamination due to autofluorescence of lipofuscin granules, which are prevalent in the GL, by ensuring that counted neurons were only present in the CR or TH channel and not in the unstained channel. Neuronal density was then calculated for each layer. CR+ neuron density was then normalized to the mean density in 7-day saline-treated mice.

We measured PV+ neuron soma area only in the EPL, where PV+ neuron density ^32^ is high enough to enable consistent sampling. We drew polygonal regions of interest (ROIs) around a total of 20 randomly selected PV+ neurons per OB section (five each in the dorsal, ventral, medial and lateral EPL). This was completed for the EPL of all images used for PV+ cell density analysis.

For analysis of CR+EdU+ neuron density, the GL and EPL were outlined, and their areas measured. Neurons that were both CR+ and EdU+ were marked using the multi-point tool, and their density was calculated. To determine the proportion of EdU+ cells that express CR, the total EdU density was calculated by counting the total number of EdU+ cells within the same outlined regions for the GL and EPL. The background-subtracted mean fluorescence intensity of CR staining was also determined for the GL and EPL in these images.

### Modeling of PV+ neuron excitability

We used a leaky integrate and fire model in Matlab 2018b (Mathworks) to determine the number of action potentials elicited by 300 ms square depolarizing current pulses (200 – 400 pA). Parameters for PV+ neurons were either stated in, or derived from action potential firing traces provided in, Kato et al. 2013^32^: resting membrane potential -60 mV; spike threshold -35 mV; post-spike voltage -60 mV; membrane time constant (tau) 5.9 ms; and input resistance 91 MΩ (for saline-treated mice). As soma area was very similar between 7- and 14-day post-MMZ groups, we used to the soma area of 7-day post-MMZ PV+ neurons, which resulted in an input resistance of 121 MΩ. The difference in input resistance was calculated from the difference in soma area between PV+ neurons from saline- and MMZ-treated mice given that tau = input resistance x capacitance, input resistance = membrane resistance/membrane surface area and capacitance = membrane capacitance x membrane surface area. The model assumes no difference in the dendritic arbors of PV+ neurons in saline vs. MMZ-treated mice. Rheobase was defined as the minimum current that elicited action potential firing. We note that the model captures the non-adapting high frequency firing reported for OB PV+ neurons but does not fully recapitulate their firing pattern in terms of a gap following initial spikes before high frequency firing resumes^32^.

### Statistics

Statistical analysis was performed using Prism 10 (GraphPad). All data sets were compared using 2-way ANOVA and Sidak’s multiple comparisons tests.

## Results

### Repopulation of OSNs in the OE five weeks after MMZ treatment

A single dose of MMZ has previously been shown to selectively ablate virtually all OSNs in the OE^24,27,52–54^. OSN neurogenesis from the preserved population of both globose basal cells and the normally quiescent horizontal basal cells then repopulates OSNs within four weeks^21,24,55,56^. Based on this, we selected time points 7, 14 and 35 days after MMZ treatment to study changes in OB interneuron populations as OSNs are lost and then repopulate. We first assessed the extent of OSN repopulation at these time points (Fig.1). We found that OSNs were ablated by MMZ treatment (Fig. 1b), with GAP43+ OSNs returning by 7 days post-MMZ (Fig. 1b,c) and an increased number of GAP43+ and some OMP+ OSNs present at 14 days post-MMZ (Fig. 1d). By 35 days post-MMZ, significant repopulation of both immature and mature OSNs had occurred (Fig. 1a,e). We concluded that MMZ treatment provides a model of degeneration and subsequent regeneration of OSNs, and hence sensory input to the OB, that would enable us to study the impact of these changes on OB interneuron populations.

**Figure 1.**
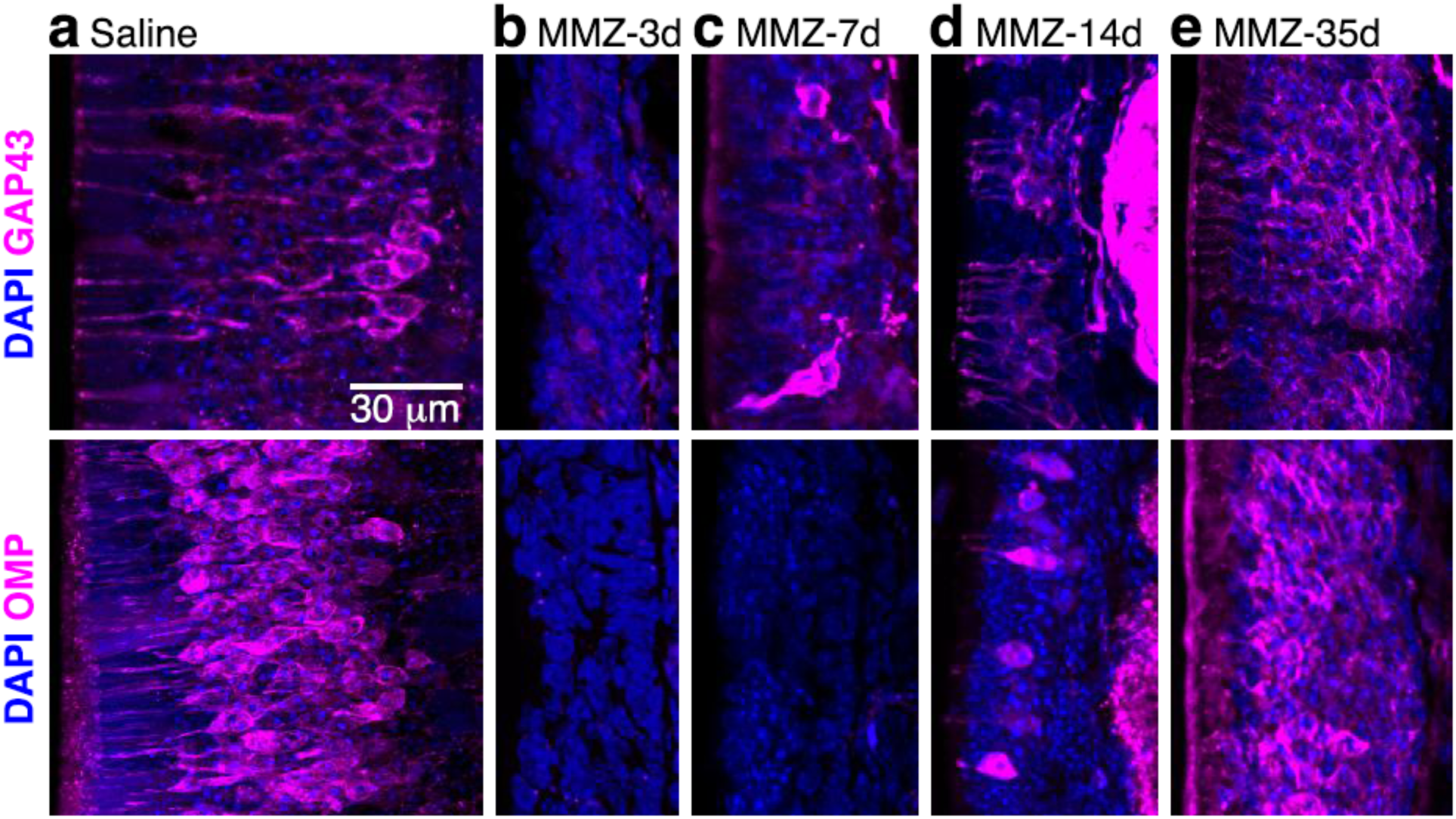
Repopulation of immature and mature OSNs after MMZ-mediated ablation. Images show single confocal optical sections of the dorsal olfactory epithelium of saline- and MMZ-treated mice. Top row: GAP43 staining of immature OSNs. Bottom row: OMP staining of mature OSNs. **a.** 3days post-saline treatment. **b-e.** 3d (**b**), 7 days (**c**), 14 days (**d**) and 35 days (**e**) post-MMZ treatment.

### Recovery of TH+ neuron density five weeks after MMZ treatment

We first assessed changes in the density of OB neurons expressing tyrosine hydroxylase (TH), the rate limiting enzyme in dopamine biosynthesis and a marker for one of the three major subtypes of inhibitory juxtaglomerular neurons in mice^57,58^. TH expression is known to be strongly dependent on sensory input in OB dopaminergic neurons^15,16,59,60^. As reported previously^15,61^, we observed a high density of TH+ neurons in the GL as well as a lower density in the EPL (Fig. 2). Therefore, we focused our analysis on those two layers. TH+ neuron density was significantly reduced in both the GL and EPL at 7 and 14 days after MMZ injection relative to saline controls (Fig. 2b-e; Fig. 3a-d). However, by 35 days after MMZ treatment, TH+ neuron density in both the GL and the EPL had recovered and was no longer significantly different to that in saline-treated mice (Fig. 2f,g; Fig. 3e,f). We found no effect of position along the anterior-posterior axis on the pattern of MMZ-related changes (Fig. 3).

**Figure 2.**
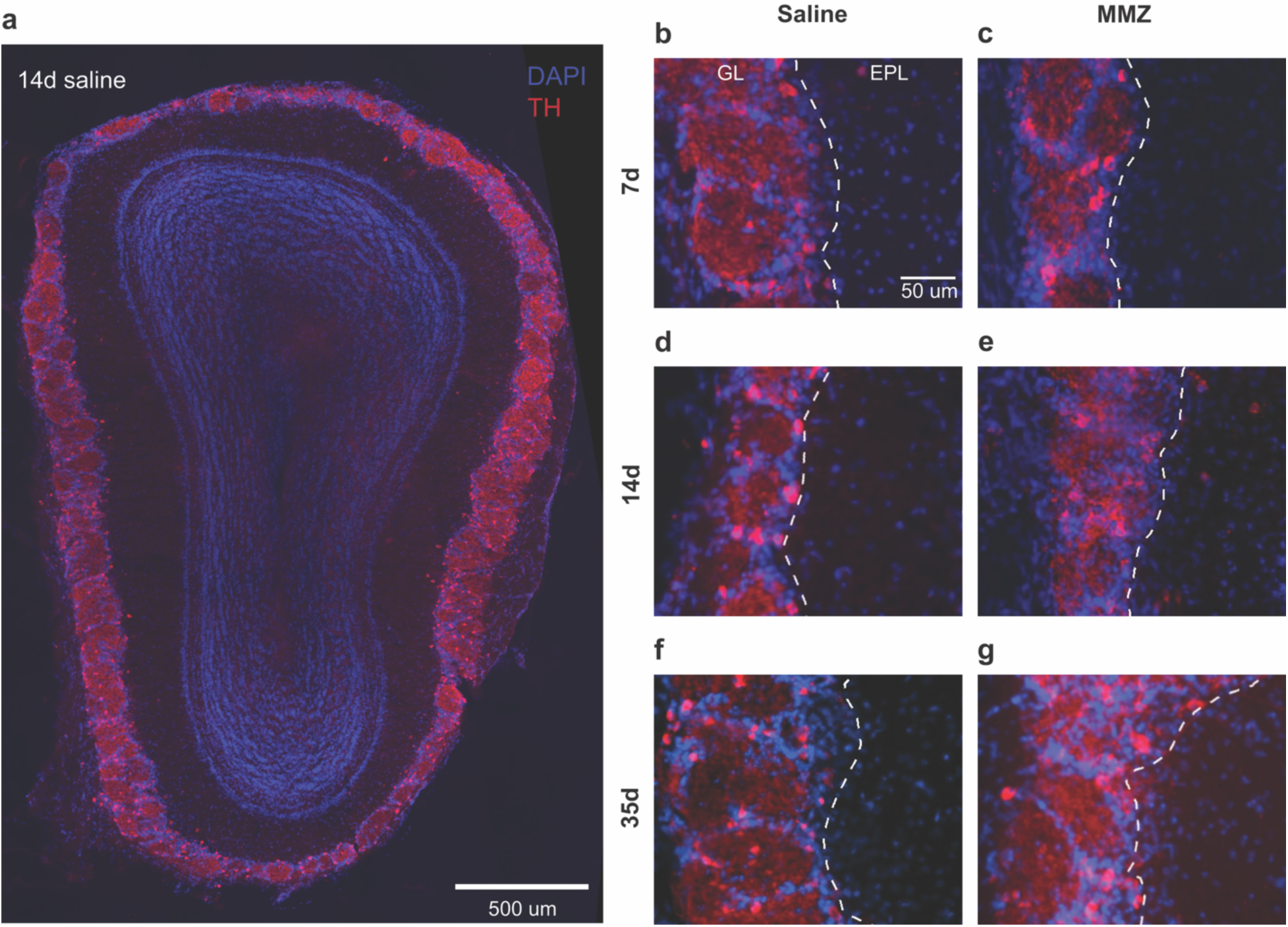
TH expression in the OB after MMZ treatment a. Coronal OB section from a saline-treated mouse showing TH expression primarily in the GL with scattered TH+ neurons in the EPL. **b-g.** Enlarged views of the ventral GL and EPL from OBs stained 7, 14 or 35 days after treatment with saline (b,d,f) or MMZ (c,e,g).

**Figure 3.**
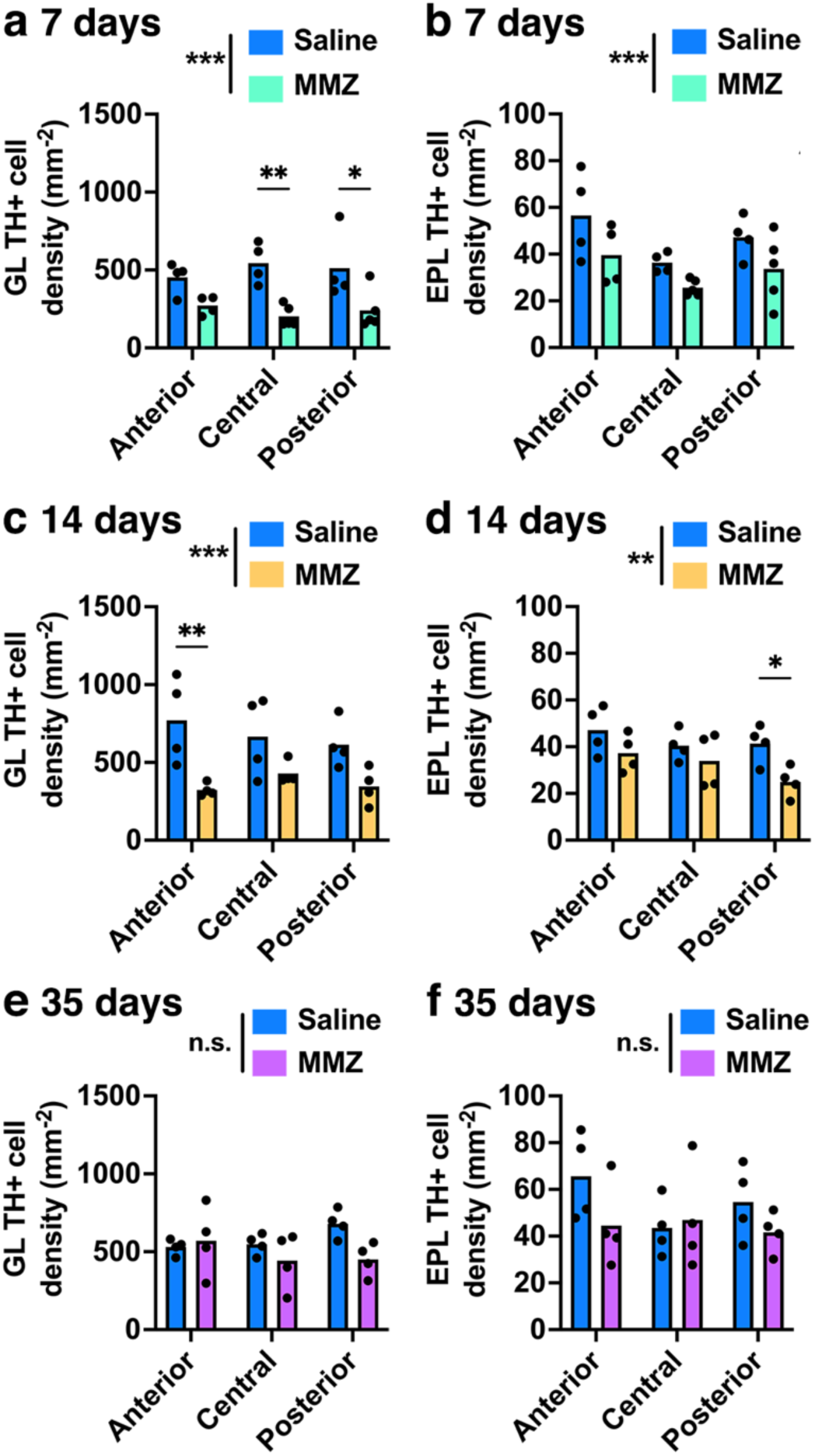
TH+ neuron density is reduced after MMZ treatment but recovers five weeks later a. GL TH+ neuron density is reduced 7 days after MMZ treatment (2-way ANOVA: P < 0.001 F_1,20_ = 27.8 for MMZ treatment, P = 0.97 F_2,20_ = 0.028 for A-P axis, P = 0.45 F_2,20_ = 0.83 for interaction). Pairwise comparisons showed a significant decrease after MMZ treatment in central and posterior sections (anterior P = 0.16, t = 2.03; central P = 0.002, t = 4.00; posterior P = 0.014, t = 3.17, Sidak’s multiple comparisons). **b.** EPL TH+ neuron density is reduced 7 days after MMZ treatment (2-way ANOVA: P = 0.008, F_1,20_ = 8.82 for MMZ treatment, P = 0.024, F_2,20_ = 4.49 for A-P axis, P = 0.86, F_2,20_ = 0.15 for interaction). **c.** GL TH+ neuron density remains reduced at 14 days after MMZ treatment (2-way ANOVA: P < 0.001 F_1,18_ = 19.5 for MMZ treatment, P = 0.68, F_2,18_ = 0.39 for A-P axis, P = 0.45, F_2,18_ = 0.83 for interaction). Pairwise comparisons showed a significant decrease after MMZ treatment in anterior sections (anterior P = 0.006, t = 3.59; central P = 0.21, t = 1.89; posterior P = 0.13, t = 2.16, Sidak’s multiple comparisons). **d.** EPL TH+ neuron density remains reduced 14 days after MMZ treatment (2-way ANOVA: P = 0.007, F_1,18_ = 9.26 for MMZ treatment, P = 0.15, F_2,18_ = 2.12 for A-P axis, P = 0.53, F_2,18_ = 0.66 for interaction). Pairwise comparisons showed a significant decrease after MMZ treatment in posterior sections (anterior P = 0.34, t = 1.58; central P = 0.67, t = 1.05; posterior P = 0.049, t = 2.64, Sidak’s multiple comparisons). **e.** GL TH+ neuron density is similar 35 days after MMZ and saline treatment (2-way ANOVA: P = 0.091, F_1,18_ = 3.19 for MMZ treatment, P = 0.56, F_2,18_ = 0.60 for A-P axis, P = 0.16, F_2,18_ = 2.02). **f.** EPL TH+ neuron density is similar 35 days after MMZ and saline treatment (2-way ANOVA: P = 0.15, F_1,18_ = 2.26 for MMZ treatment, P = 0.49, F_2,18_ = 0.74 for A-P axis, P = 0.34, F_2,18_ = 1.13 for interaction).

### MMZ treatment does not affect OB PV+ neuron density

We then asked whether the density of OB PV-expressing interneurons was affected by MMZ treatment. PV+ neurons are normally generated only during embryonic and early postnatal development^28^. However, it has been unclear whether PV+ neuron survival is affected by sensory input or whether injury can induce PV+ neurogenesis. We performed PV staining in anterior, central and posterior OB sections from mice that had been injected with MMZ or saline 7, 14 or 35 days earlier. Similar to previous reports^4,28,31,33^, we found PV+ neurons throughout the EPL, with a lower density in the MCL and IPL, and very sparsely distributed in the GCL, in both saline and MMZ-treated mice (Fig. 4). We quantified the density of PV+ neurons in these layers, treating the MCL and IPL as a single region of interest for analysis due to the limited thickness of these layers (Fig. 4b-g). We found no significant effect of either MMZ treatment or stage along the anterior-posterior axis on the density of PV+ neurons in any layer (Fig. 4,5). Therefore, we concluded that MMZ treatment does not affect PV+ neuron survival or induce PV+ neurogenesis in adult mice.

**Figure 4.**
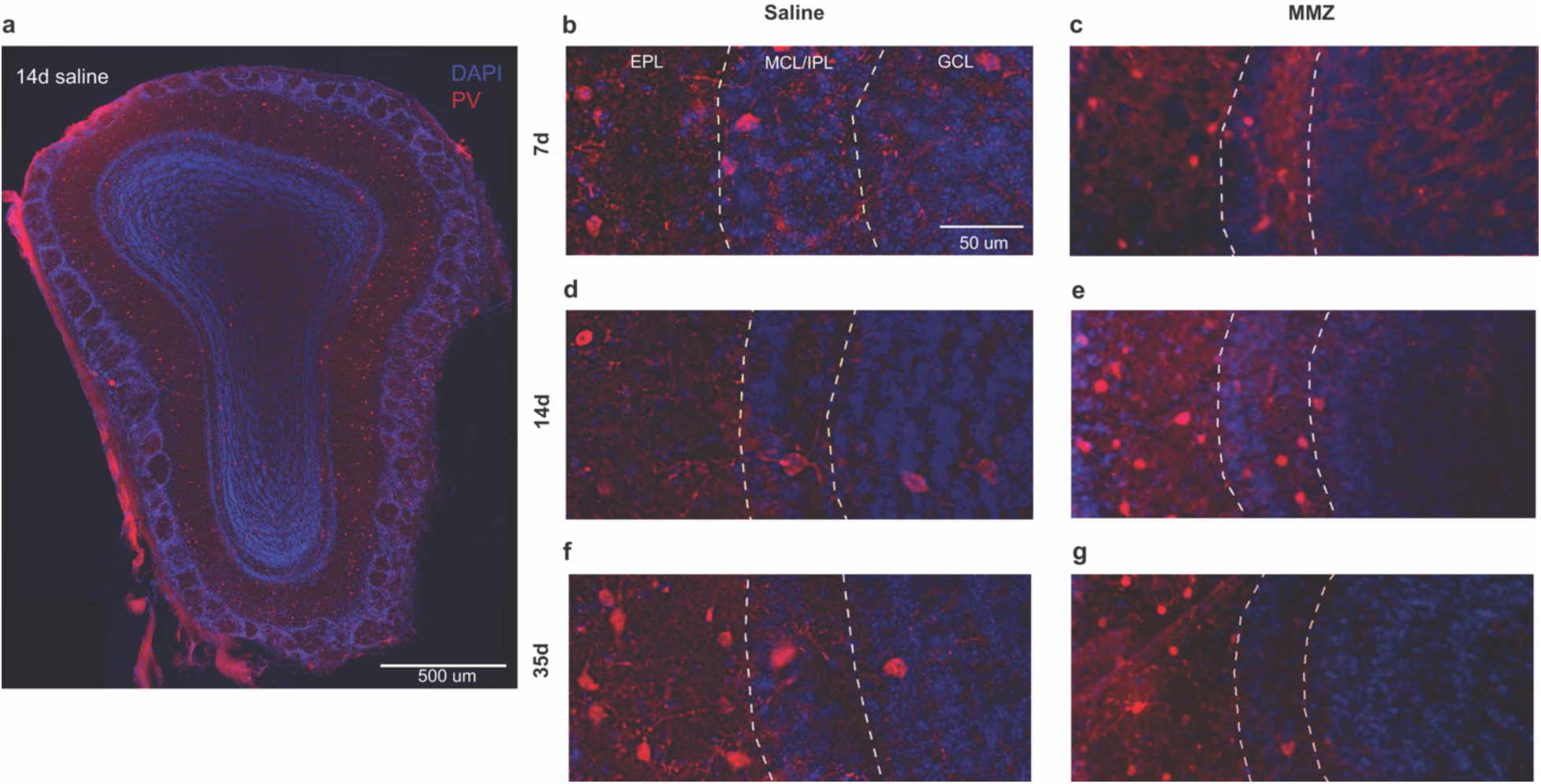
PV expression in the OB after MMZ treatment a. Coronal OB section from a saline-treated mouse showing PV expression predominantly in the EPL, with sparser expression in the MCL, IPL and GCL. **b-g.** Enlarged views of the ventral EPL, MCL, IPL and GCL from OBs stained 7, 14 or 35 days after treatment with saline (b, d, f) or MMZ (c, e, g).

**Figure 5.**
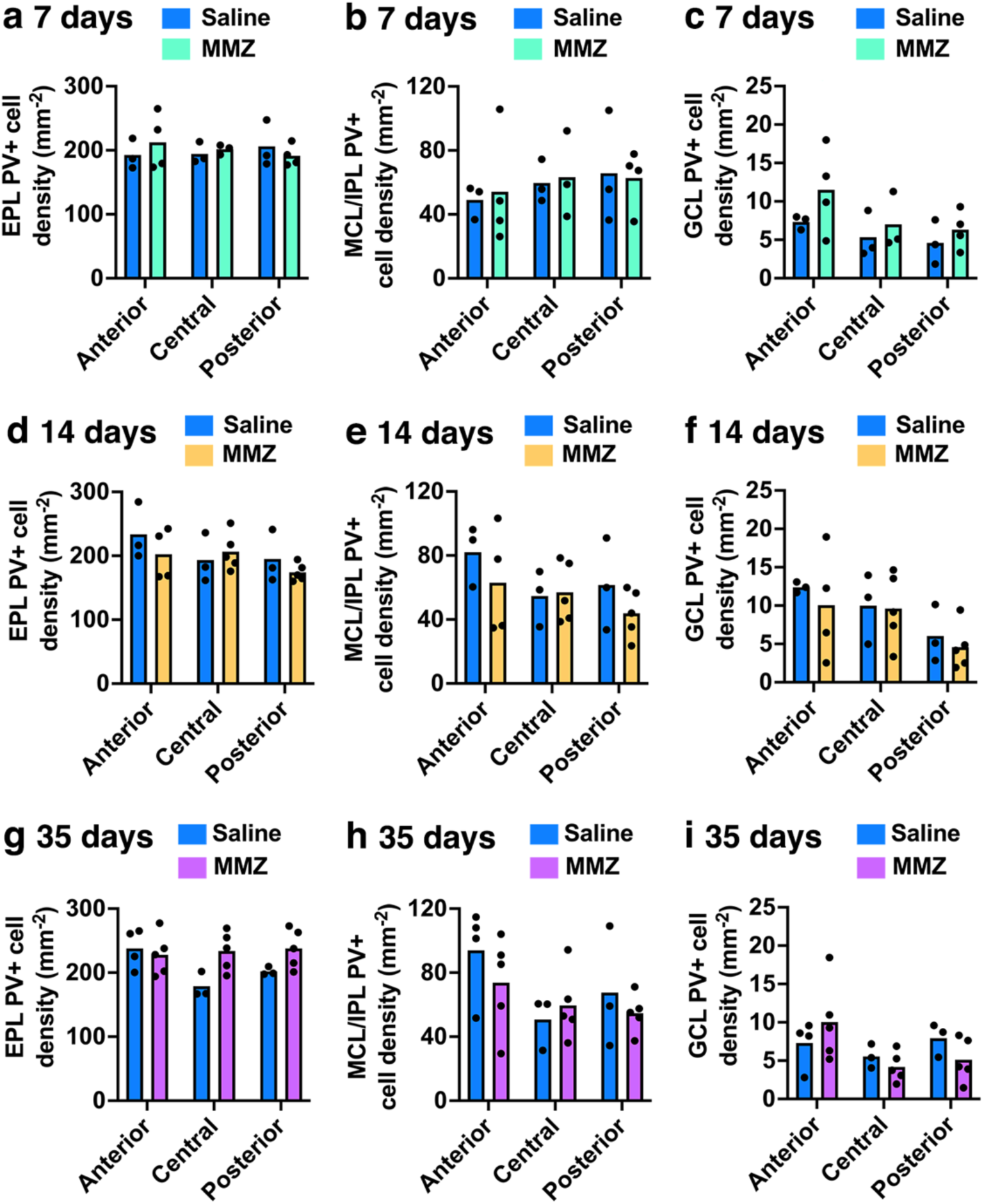
PV+ OB neuron density is not affected by MMZ treatment a. EPL PV+ neuron density is similar 7 days after MMZ and saline treatment (2-way ANOVA: P = 0.75, F_1,14_ = 0.11 for MMZ treatment, P = 0.95, F_2,14_ = 0.050 for A-P axis, P = 0.53, F_2,14_ = 0.66 for interaction). **b.** MCL and IPL PV+ neuron density is similar 7 days after MMZ and saline treatment (2-way ANOVA: P = 0.87, F_1,14_ = 0.027 for MMZ treatment, P = 0.65, F_2,14_ = 0.45 for A-P axis, P = 0.96, F_2,14_ = 0.046 for interaction). **c.** GCL PV+ neuron density is similar 7 days after MMZ and saline treatment (2-way ANOVA: P = 0.14, F_1,14_ = 2.49 for MMZ treatment, P = 0.13, F_2,14_ = 2.42 for A-P axis, P = 0.76, F_2,14_ = 0.28 for interaction). **d.** EPL PV+ neuron density is similar 14 days after MMZ and saline treatment (2-way ANOVA: P = 0.38, F_1,17_ = 0.83 for MMZ treatment, P = 0.20, F_2,17_ = 1.77 for A-P axis, P = 0.44, F_2,17_ = 0.86 for interaction). **e.** MCL and IPL PV+ neuron density is similar 14 days after MMZ and saline treatment (2-way ANOVA: P = 0.25, F_1,17_ = 1.41 for MMZ treatment, P = 0.24, F_2,17_ = 1.58 for A-P axis, P = 0.61, F_2,17_ = 0.51 for interaction). **f.** GCL PV+ neuron density is similar 14 days after MMZ and saline treatment (2-way ANOVA: P = 0.48, F_1,17_ = 0.52 for MMZ treatment, P = 0.056, F_2,17_ = 3.44 for A-P axis, P = 0.92, F_2,17_ = 0.085 for interaction). **g.** EPL PV+ neuron density is similar 35 days after MMZ and saline treatment (2-way ANOVA: P = 0.094, F_1,19_ = 3.11 for MMZ treatment, P = 0.44, F_2,19_ = 0.86 for A-P axis, P = 0.20, F_2,19_ = 1.75 for interaction). **h.** MCL and IPL PV+ neuron density is similar 35 days after MMZ and saline treatment (2-way ANOVA: P = 0.44, F_1,19_ = 0.62 for MMZ treatment, P = 0.071, F_2,19_ = 3.05 for A-P axis, P = 0.50, F_2,19_ = 0.71 for interaction). **i.** GCL PV+ neuron density is similar 35 days after MMZ and saline treatment (2-way ANOVA: P = 0.72, F_1,19_ = 0.13 for MMZ treatment, P = 0.082, F_2,19_ = 2.86 for A-P axis, P = 0.22, F_2,19_ = 1.64 for interaction).

However, we observed that the somatic morphology of PV+ neurons throughout the analyzed layers differed between MMZ- and saline-treated mice (Fig. 4b-e). To quantify this, we focused on the EPL, where PV+ neurons are numerous, and measured the soma area of a sample of PV neurons for each mouse. We found that PV+ neuron soma area was significantly reduced 7 and 14 days after MMZ treatment (Fig. 6a,b) but recovered to baseline by 35 days post-MMZ (Fig. 6c). Therefore, we concluded that OSN ablation induces activity dependent changes in OB PV+ neuron soma size. To determine the functional impact of reduced soma area, we determined the resultant change in input resistance and then used a leaky integrate and fire model to compare responses to square depolarizing current pulses in PV+ neurons. We found that rheobase was reduced and firing rate was significantly increased in the small soma area/higher input resistance PV+ neurons that are present 7-14 days after MMZ treatment (Fig. 6d). Hence, PV+ neurons are likely to have increased intrinsic excitability in 7-14-day post-MMZ mice.

**Figure 6.**
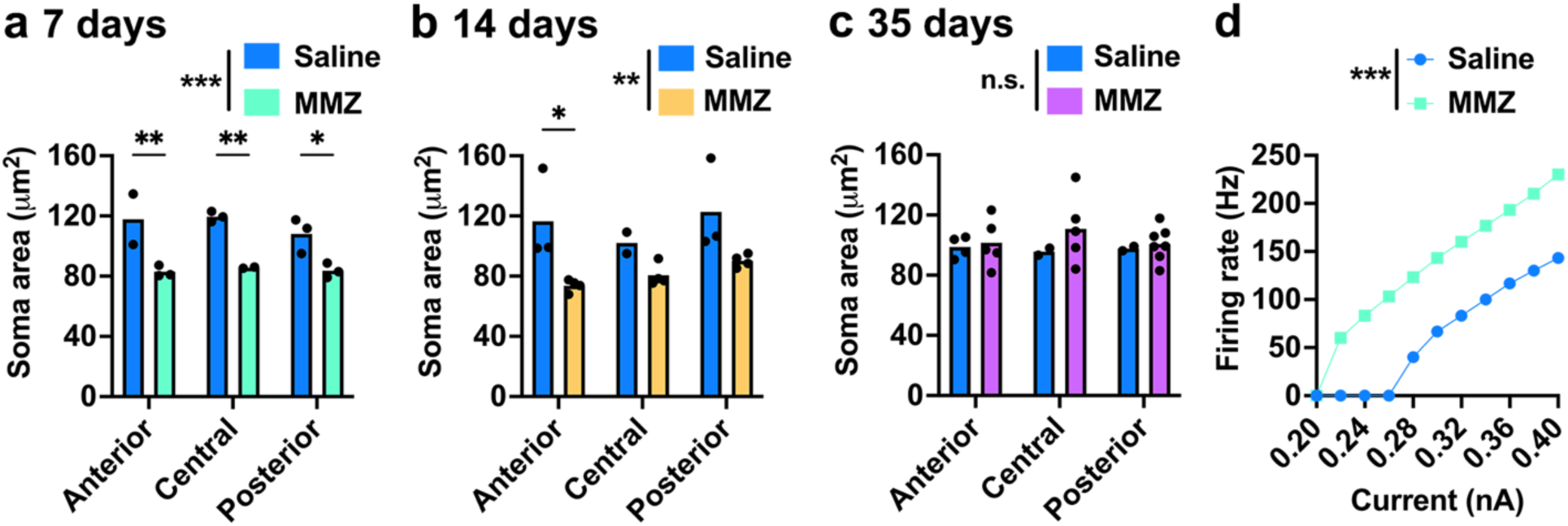
Reduced soma size of PV+ EPL neuron 7-14 days after MMZ treatment a. Soma area of EPL PV+ neurons is reduced 7 days after MMZ treatment (2-way ANOVA: P < 0.001, F_1,10_ = 39.71 for MMZ treatment, P = 0.52, F_2,10_ = 0.69 for A-P axis, P = 0.63, F_2,10_ = 0.49 for interaction). Pairwise comparisons showed a significant decrease after MMZ treatment in anterior, central and posterior sections (anterior P = 0.034, t = 3.95; central P = 0.010, t = 3.83; posterior P = 0.008, t = 3.09, Sidak’s multiple comparisons). **b.** Soma area of EPL PV+ neurons remains reduced 14 days after MMZ treatment (2-way ANOVA: P = 0.001, F_1,14_ = 16.49 for MMZ treatment, P = 0.30, F_2,14_ = 1.32 for A-P axis, P = 0.58, F_2,14_ = 0.57 for interaction). Pairwise comparisons showed a significant decrease after MMZ treatment in anterior sections (anterior P = 0.017, t = 3.25; central P = 0.43, t = 1.44; posterior P = 0.078, t = 2.47, Sidak’s multiple comparisons). **c.** Soma area of EPL PV+ neurons is similar 35 days after MMZ and saline treatment (2-way ANOVA: P = 0.29, F_1,19_ = 1.17 for MMZ treatment, P = 0.89, F_2,19_ = 0.12 for A-P axis, P = 0.70, F_2,19_ = 0.37 for interaction). **d.** Predicted firing rate of PV+ neurons is higher in 7-day MMZ vs. 7-day saline treated mice from a leaky integrate and fire model (paired t-test: P < 0.001, t = 9.21).

### Transient increase in CR+ neuron density in the GL and EPL 7 days after MMZ treatment

The effects of MMZ treatment on CR+ neuron survival or neurogenesis are unknown. In saline-treated mice, we found a high density of CR+ neurons in the GL and superficial GCL, a lower density in the EPL, and only scattered neurons in the MCL and IPL (Fig. 7), similar to previous reports^43,62,63^. We therefore focused our quantitative analysis on CR-expressing neurons in the GL, EPL and GCL.

**Figure 7.**
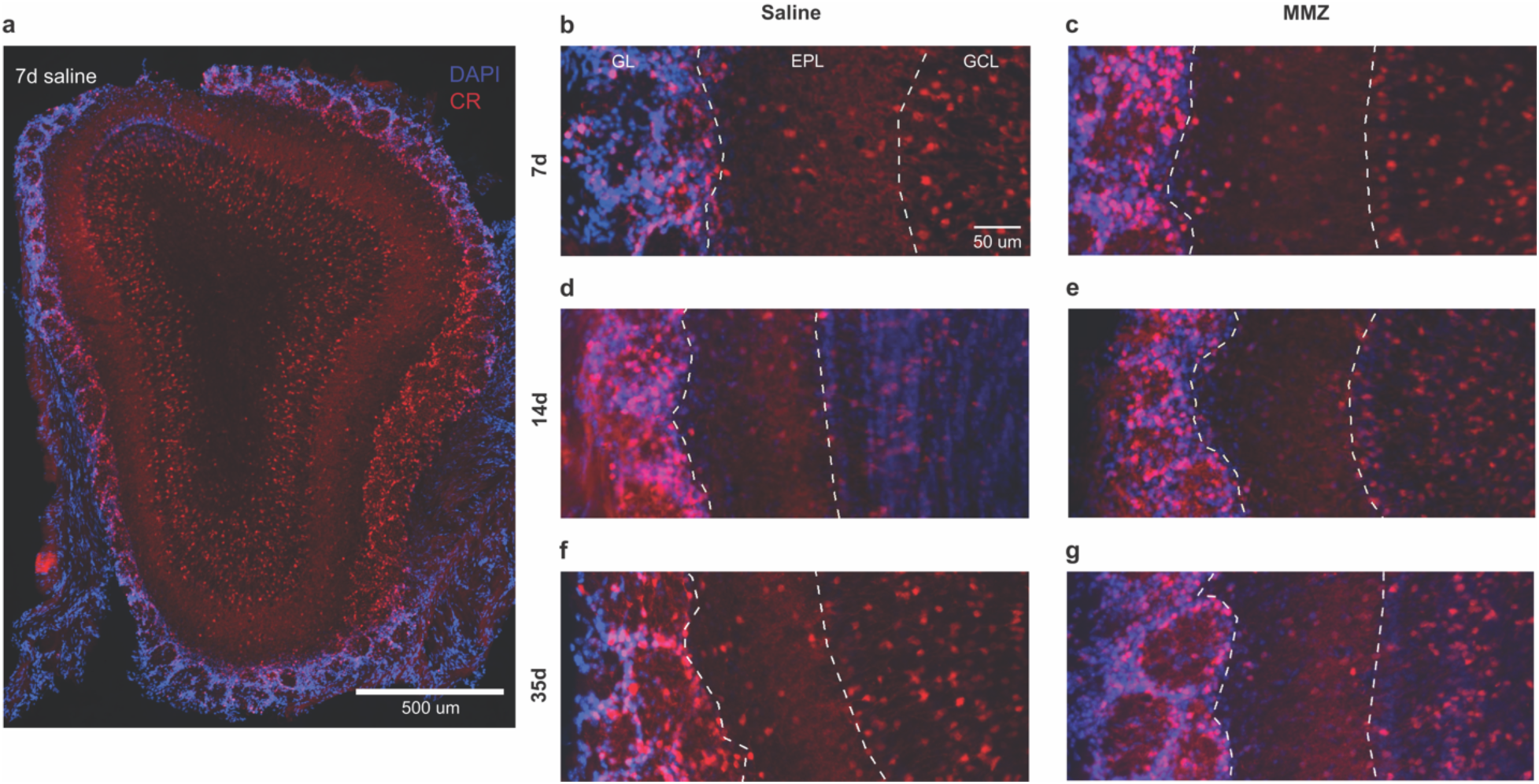
CR expression in the OB after MMZ treatment a. Coronal OB section from a saline-treated mouse showing CR expression throughout the GL, EPL and GCL. **b-g.** Enlarged views of the ventral GL, EPL and GCL from OBs stained 7, 14 or 35 days after treatment with saline (b, d, f) or MMZ (c, e, g).

At the 7-day time point, we found that the density of CR+ neurons in the GL and EPL was consistently significantly higher in MMZ-treated than in saline-treated mice, throughout the anterior-posterior axis (Fig. 7b,c; Fig. 8a,b). In contrast, CR+ neuron density in the GCL was similar in MMZ- and saline-treated mice at the 7-day time point (Fig. 8c). At the 14-day time point, there was no longer a difference in CR+ neuron density in the GL or EPL between MMZ- and saline-treated mice (Fig. 7d,e; Fig. 8d-e), but GCL CR+ neuron density was lower in MMZ-treated mice (Fig. 8f). In pairwise comparisons of sections at different stages through the OB, GCL CR+ neuron density was only significantly lower in MMZ-treated mice for anterior sections, although central and posterior sections showed a similar trend. At the 35-day time point, there was no difference in CR+ neuron density in any layer between MMZ- and saline-treated mice (Fig. 7f,g; Fig. 8g-i).

**Figure 8.**
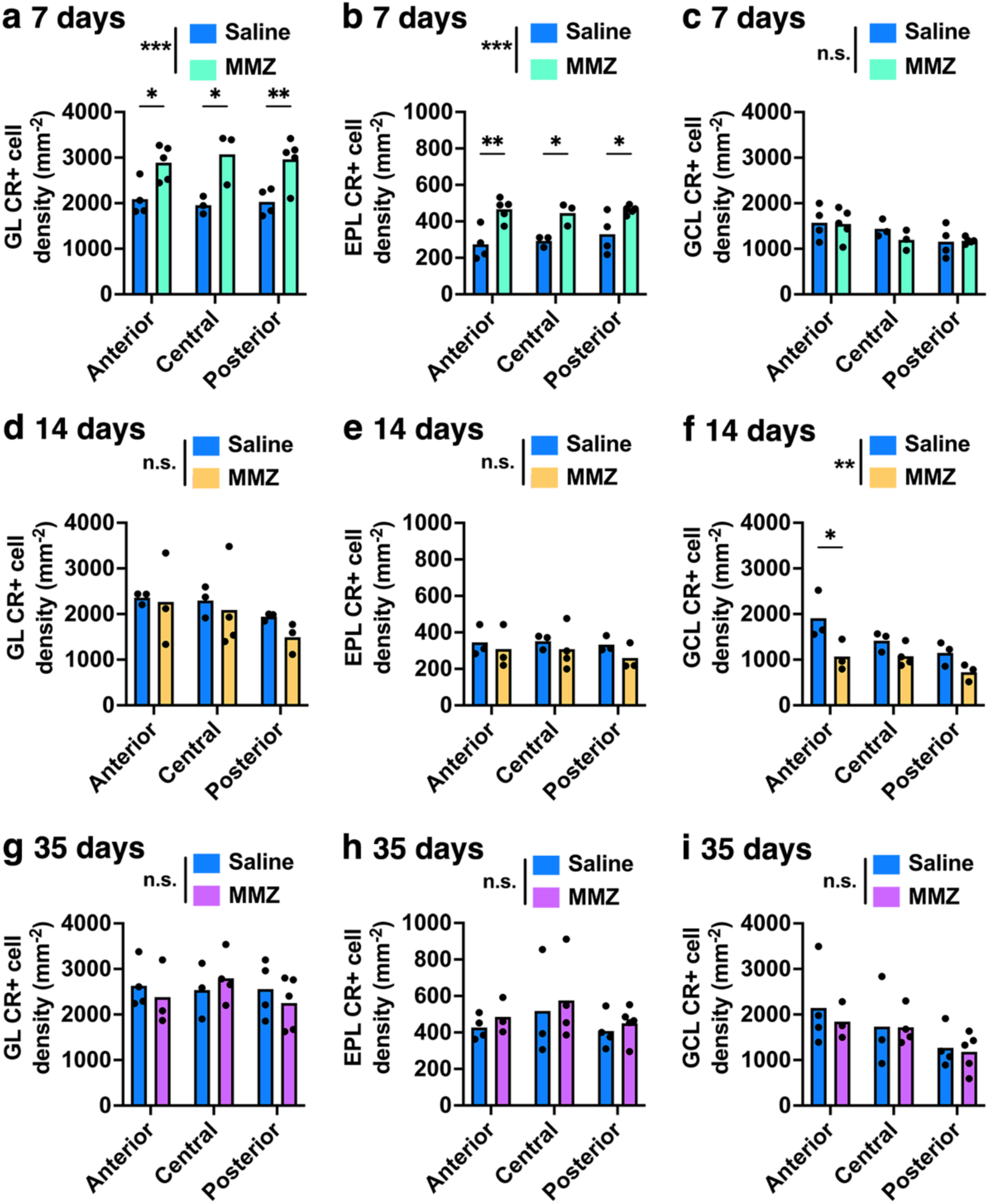
Transient increase in CR+ neuron density in the GL and EPL 7 days after MMZ treatment a. GL CR+ neuron density is increased 7 days after MMZ treatment (2-way ANOVA: P < 0.001, F_1,18_ = 30.62 for MMZ treatment, P = 0.99, F_2,18_ = 0.006 for A-P axis, P = 0.77, F_2,18_ = 0.27 for interaction). Pairwise comparisons showed a significant increase after MMZ treatment in anterior, central and posterior sections (anterior P = 0.029, t = 2.89; central P = 0.011, t = 3.33; posterior P = 0.010, t = 3.38, Sidak’s multiple comparisons). **b.** EPL CR+ neuron density is increased 7 days after MMZ treatment (2-way ANOVA: P < 0.001, F_1,18_ = 30.42 for MMZ treatment, P = 0.67, F_2,18_ = 0.42 for A-P axis, P = 0.67, F_2,18_ = 0.40 for interaction). Pairwise comparisons showed a significant increase after MMZ treatment in anterior, central and posterior sections (anterior P = 0.002, t = 4.13; central P = 0.043, t = 2.70; posterior P = 0.030, t = 2.88, Sidak’s multiple comparisons). **c.** GCL CR+ neuron density is similar 7 days after MMZ and saline treatment but there was a significant effect of position along the anterior-posterior axis (2-way ANOVA: P = 0.49, F_1,18_ = 0.50 for MMZ treatment, P = 0.029, F_2,18_ = 4.35 for A-P axis, P = 0.66, F_2,18_ = 0.43 for interaction). Pairwise comparisons showed that CR+ neuron density was lower in posterior than in anterior sections (P = 0.027, t = 2.93, Sidak’s multiple comparisons). **d.** GL CR+ neuron density is similar 14 days after MMZ and saline treatment (2-way ANOVA: P = 0.41, F_1,13_ = 0.72 for MMZ treatment, P = 0.27, F_2,13_ = 1.47 for A-P axis, P = 0.89, F_2,13_ = 0.12 for interaction). **e.** EPL CR+ neuron density is similar 14 days after MMZ and saline treatment (2-way ANOVA: P = 0.23, F_1,13_ = 1.57 for MMZ treatment, P = 0.77, F_2,13_ = 0.27 for A-P axis, P = 0.93, F_2,13_ = 0.072 for interaction). **f.** GCL CR+ neuron density is reduced 14 days after MMZ treatment (2-way ANOVA: P = 0.003, F_1,13_ = 13.60 for MMZ treatment, P = 0.030, F_2,13_ = 4.68 for A-P axis, P = 0.35, F_2,13_ = 1.15 for interaction). Pairwise comparisons showed a significant decrease after MMZ treatment in anterior sections (anterior P = 0.018, t = 3.29; central P = 0.45, t = 1.41; posterior P = 0.33, t = 1.65, Sidak’s multiple comparisons), and that CR+ neuron was density was lower in posterior than in anterior sections (P = 0.028, t = 3.05, Sidak’s multiple comparisons). **g.** GL CR+ neuron density is similar 35 days after MMZ and saline treatment (2-way ANOVA: P = 0.70, F_1,17_ = 0.15 for MMZ treatment, P = 0.69, F_2,17_ = 0.37 for A-P axis, P = 0.62, F_2,17_ = 0.49 for interaction). **h.** EPL CR+ neuron density is similar 35 days after MMZ and saline treatment (2-way ANOVA: P = 0.45, F_1,17_ = 0.60 for MMZ treatment, P = 0.36, F_2,17_ = 1.10 for A-P axis, P = 0.99, F_2,17_ = 0.006 for interaction). **i.** GCL CR+ neuron density is similar 35 days after MMZ and saline treatment (2-way ANOVA: P = 0.62, F_1,17_ = 0.26 for MMZ treatment, P = 0.072, F_2,17_ = 3.09 for A-P axis, P = 0.91, F_2,17_ = 0.097 for interaction).

We also found differences in GCL CR+ neuron density along the anterior-posterior axis: density was higher in anterior than in posterior sections at both the 7-day (Fig. 8c) and the 14-day (Fig. 8f) time points, and there was a similar trend that did not reach statistical significance at the 35-day time point (Fig. 8i). To determine whether GCL CR+ neuron density may be non-uniform in the healthy OB, we further analyzed our data from saline-injected mice. We found a significant effect of anterior-posterior axis position (P = 0.031, F_2,22_ = 4.09) but not of time point after saline treatment (P = 0.39, F_2,22_ = 0.98; interaction P = 0.93, F_4,22_ = 0.22, 2-way ANOVA). Pairwise comparisons showed that GCL CR+ neuron density was higher in anterior than in posterior sections (P = 0.024, q = 4.04, Tukey’s multiple comparisons) but found no difference between anterior vs. central or central vs. posterior sections. Therefore, our data suggest that there is a subtle gradient in GCL CR+ neuron density along the anterior-posterior axis of the OB.

To better understand the temporal progression of these changes, for each mouse, we calculated the CR+ neuron density in each layer, pooling data from sections along the anterior-posterior axis. We then normalized the CR density values for each mouse to the mean of the 7-day saline-treated group. We found that in MMZ-treated mice, the density of CR+ neurons decreased significantly between the 7- and 14-day time points in the GL (Fig. 9a) and showed a similar trend in the EPL that did not reach statistical significance (Fig. 9b). While CR+ neuron density was stable between the 14- and 35-day time points in the GL, it increased significantly in both the EPL and the GCL between these time points (Fig. 9a-c). Therefore, MMZ treatment may cause an upregulation of neurogenesis that increases CR+ neuron density in the EPL and GCL.

**Figure 9.**
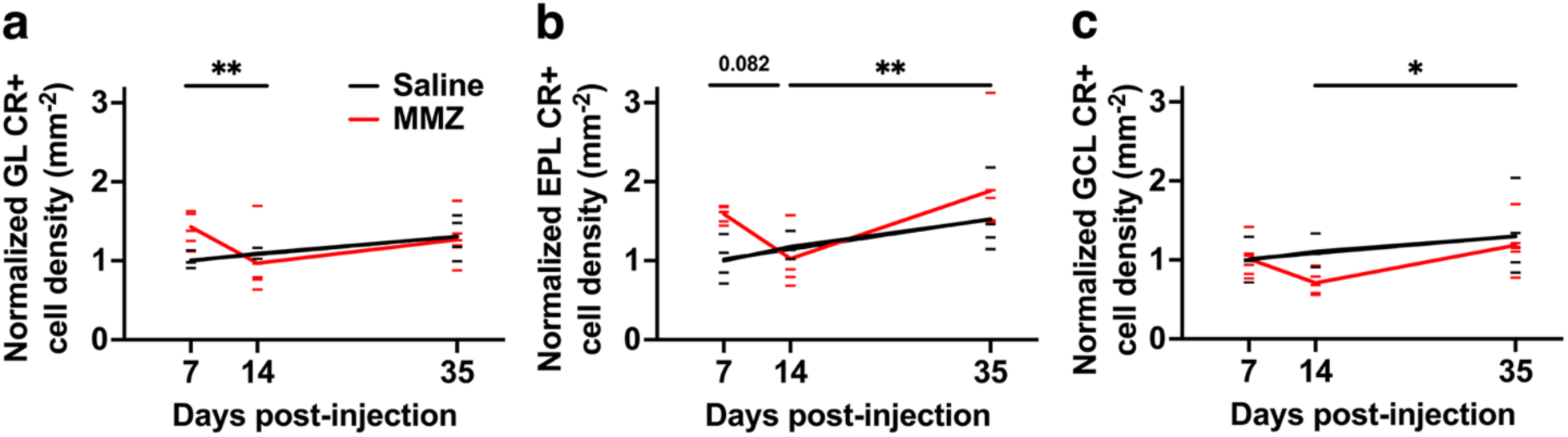
MMZ-dependent changes in normalized CR+ neuron density a. GL normalized CR+ neuron density in the GL decreased between 7 and 14d post-MMZ (7d vs. 14d MMZ, P = 0.031, t = 2.81; all other comparisons, P > 0.22; Sidak’s multiple comparisons test after 2-way ANOVA**). b.** EPL normalized CR+ neuron density showed a trend to decrease between 7d and 14d post-MMZ and increased between 14d and 35d in MMZ-treated mice (7d vs. 14d MMZ, P = 0.082, t = 2.35; 14d vs. 35d MMZ, P = 0.005, t = 3.57; all other comparisons, P > 0.22; Sidak’s multiple comparisons test after 2-way ANOVA). **c.** GCL normalized CR+ neuron density increased between 14d and 35d in MMZ-treated mice (14d vs. 35d MMZ, P = 0.042, t = 2.66; all other comparisons, P > 0.28, Sidak’s multiple comparisons test after 2-way ANOVA).

Pairwise comparisons did not reveal any differences in CR+ neuron density between time points in saline-treated mice in any layer; however, we noticed that all three layers showed a trend for CR+ neuron density to increase over time (Fig. 9). We therefore performed linear regression analysis for CR+ neuron density in each layer in saline-treated mice. We found a significant positive relationship between time and CR+ neuron density in the GL (R^2^ = 0.43, P = 0.028, F_1,9_ = 6.88) and the EPL (R^2^ = 0.37, P = 0.046, F_1,9_ = 5.33), but no significant relationship in the GCL (R^2^ = 0.14, P = 0.27, F_1,9_ = 1.41). We therefore concluded that ongoing neurogenesis may increase CR+ neuron density across the four-week span of our experimental time points, at least in the GL and EPL.

We next focused on potential mechanisms that could underlie the transient increase in GL and EPL CR+ neuron density 7 days after MMZ treatment. CR+ neurons throughout the OB continue to be generated in adult mice^28,35^; therefore, we asked whether enhanced CR+ neurogenesis could underlie the increase in CR+ neuron density that we observed. Approximately 90% of new juxtaglomerular neurons generated in the SVZ take over 7 days to reach the GL^64^, making it unlikely that upregulation of SVZ neurogenesis in response to MMZ-induced loss of sensory input could underlie the rapid increase in GL and EPL CR+ neuron density. However, it was possible that neurogenesis in the core of the OB itself, which preferentially produces CR+ neurons^37,39–41^, could enable newly generated CR+ neurons to reach the GL and EPL within 7 days of MMZ injection. Therefore, we injected mice with either MMZ or saline, and then one day later with EdU, a thymidine analog that labels terminally dividing cells. We then quantified newborn CR+ neurons in the GL and EPL (Fig. 10a-d).

**Figure 10.**
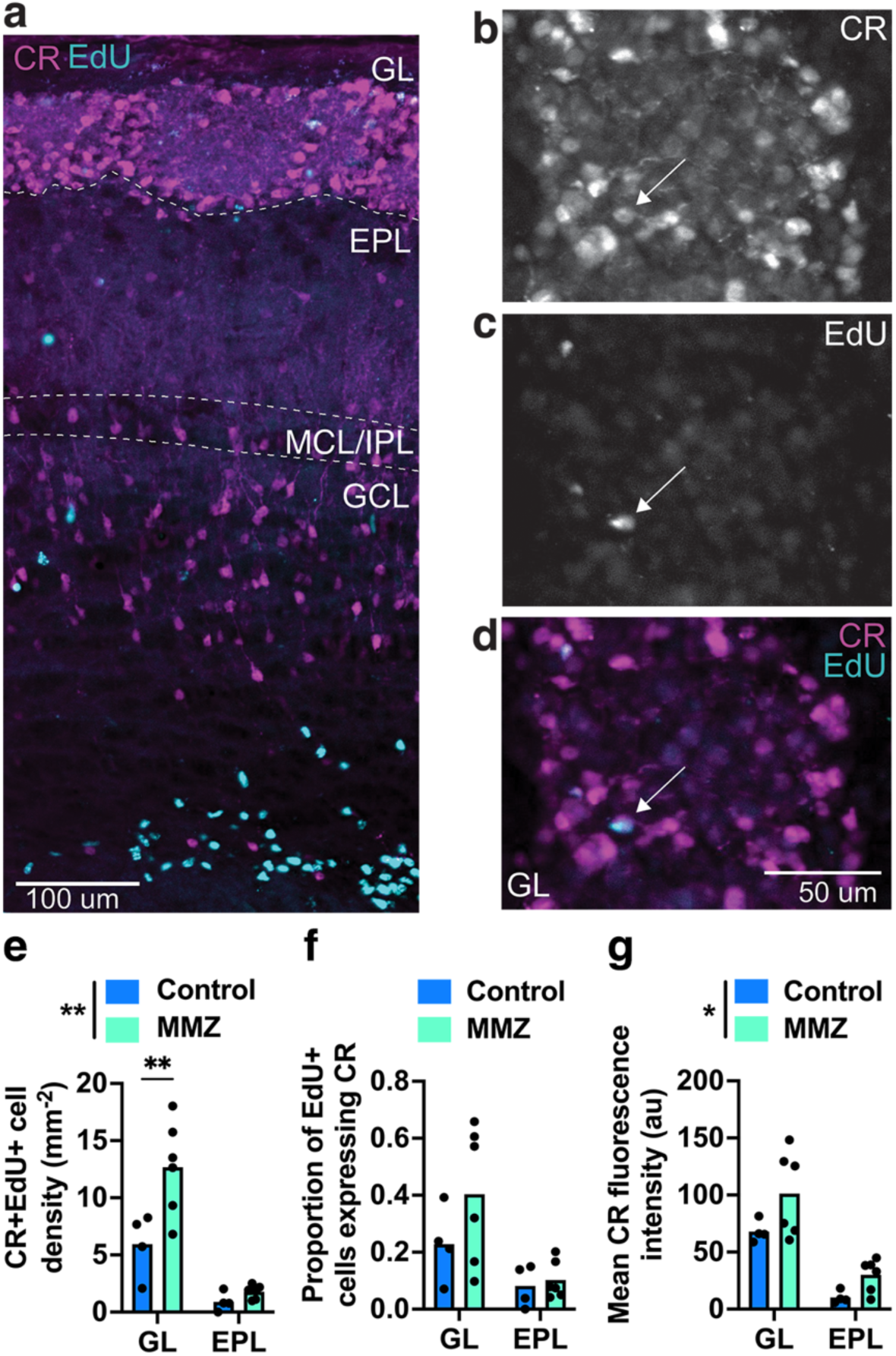
Increased CR expression rather than neurogenesis can account for the MMZ-mediated increase in CR+ neuron density a. Coronal OB section from a MMZ-treated mouse showing CR expression and EdU labeling throughout the OB 7 days after MMZ and 6 days after EdU treatment. **b-d.** Enlarged view of a glomerulus showing CR expression (b), EdU labeling (c) and colocalization of CR and EdU (d, arrow). **e.** GL and EPL CR+EdU+ neuron density is increased 7 days after MMZ treatment (2-way ANOVA: P = 0.006, F_1,16_ = 9.96 for MMZ treatment, P < 0.001, F_1,16_ = 43.69 for layer, P = 0.027, F_1,16_ = 5.92 for interaction). Pairwise comparisons showed a significant increase after MMZ treatment in the GL (GL P = 0.002, t = 3.95; EPL P = 0.85, t = 0.51, Sidak’s multiple comparisons). **f.** Proportion of EdU+ cells that express CR in the GL and EPL is not affected by MMZ treatment in the GL or EPL (2-way ANOVA: P = 0.18, F_1,16_ = 1.92 for MMZ treatment, P = 0.006, F_1,16_ = 10.1 for layer, P = 0.29, F_1,16_ = 1.21 for interaction). **g.** GL and EPL CR fluorescence intensity is increased 7 days after MMZ treatment (2-way ANOVA: P = 0.021, F_1,16_ = 6.50 for MMZ treatment, P < 0.001, F_1,16_ = 38.28 for layer, P = 0.53, F_1,16_ = 0.42 for interaction).

We found a two-fold increase in the density of CR+EdU+ neurons in both the GL and the EPL in MMZ- vs. saline-treated mice (Fig. 10e). However, even in MMZ-treated mice, the density of CR+EdU+ neurons was very low compared to the increase in CR+ neuron density due to MMZ treatment (GL: 6 mm^-2^ CR+EdU+; 861 mm^-2^ increase in CR+ neuron density 7-days post-MMZ), and hence could not explain the overall increase in CR+ neuron density. We next determined the proportion of EdU-labeled cells that expressed CR but found no difference between MMZ-treated and control mice (Fig. 10f). Finally, we considered whether an increase in CR expression in existing GL and EPL neurons could underlie the increased CR+ neuron density that we observed. To estimate CR expression, we quantified the mean fluorescence intensity of CR staining in the GL and EPL, in images collected using identical acquisition settings. We found a significant increase in mean CR fluorescence intensity in the GL and EPL of MMZ-treated mice (Fig. 10g), with a 49% increase in the GL and a 192% increase in the EPL. We concluded that CR expression is increased 7d after MMZ treatment.

## Discussion

### Methimazole treatment as a model for loss and regeneration of OB sensory input

In agreement with previous studies^24–27,52,65,66^, following MMZ treatment, we found complete ablation of OSNs within 3 days, with newly generated immature OSNs present within 7 days, newly generated mature OSNs present by 14 days, and substantial OSN repopulation by 35 days. There is also good evidence for recovery of sensory input to the OB. We have shown previously that newly generated, immature OSN axons reach the OB within 5 days of MMZ treatment, and that odor detection and discrimination behavior begins to recover within a week^27^. There is also good evidence for recovery of OB sensory input even after treatment with 100 mg/kg MMZ (vs. 75 mg/kg used in this study). OSN connectivity with external tufted cells in the OB recovered to control levels within 16 days, with maturation of presynaptic release properties complete within a further week^25^. Furthermore, OSN projections to the M72 and I7 glomeruli, and performance in a previously learnt odor discrimination task, recovered within 45 days of MMZ treatment^26^. The pattern of changes in TH+ neuron density that we saw also supports a dramatic reduction in sensory input, followed by gradual regeneration of sensory input between 7 and 35 days post-MMZ. It is highly likely that the reduction in TH+ neuron density in the GL and EPL arises from a combination of reduced TH expression and reduced DA neuron survival^14–19^. Similarly, both activity-dependent recovery of TH expression and addition of newborn DA neurons are likely to contribute to the substantial recovery of TH+ neuron density by 35 days post-MMZ^15,18,19^. Overall, we concluded that MMZ treatment enables the impact of bidirectional changes in OB sensory input to be assessed.

### Reduced soma size but unchanged density of PV+ neurons after MMZ treatment

PV+ neurons are normally generated predominantly during mouse embryonic development, with neurogenesis complete before P30^28^. We therefore investigated PV+ neuron density to determine whether MMZ can reactivate neurogenesis of OB interneuron subtypes that are not normally generated during adulthood. However, we found no effect of MMZ treatment on PV+ neuron density in any OB layer at any time point examined. This indicates both that PV expression in the OB is not activity-dependent, and that generation of PV+ neurons in the SVZ is not reactivated following MMZ treatment in mice. This may suggest more broadly that certain subtypes of SVZ-generated neurons cannot be produced in the adult brain, irrespective of experimental manipulations. There is considerable regional heterogeneity amongst SVZ NSCs, arising both from embryonic regional specification and differential transcription factor expression^67,68^. PV+ neurons are generated from neural stem cells in both the pallial and lateral/medial ganglionic eminence regions of the telencephalic neuroepithelium (and the dorsal and lateral wall regions derived from them in the early postnatal SVZ)^69^. However, these same SVZ regions continue to generate new CR+ and TH+ OB interneurons in adult mice^69^, suggesting a finer level of control over developmental differences in interneuron subtype-specific neurogenesis.

Our findings differ from those of a previous study that found reduced PV+ neuron density following naris occlusion beginning either perinatally or at P30 in the EPL of rats^4^. The authors attributed this to an activity-dependent reduction in PV expression, for which there is precedent in rat binocular visual cortex following monocular deprivation^70^, although they did not rule out cell death. Notably, PV+ neuron density is much lower in rats (∼10 neurons mm^-2^ ^4^) than in mice (∼200 neurons mm^-2^, Fig. 6), and PV+ neurogenesis also peaks later in rats^4,28,29^, making it unlikely that PV+ neurons in rats make the same contribution to mitral and tufted cell inhibition as those in mice^32^. Given the morphological heterogeneity of PV+ neurons in mice^34^, it is plausible that rat EPL PV+ neurons correspond to a quantitatively minor PV+ neuron subtype in mice, with changes in the density of that subtype undetectable against the backdrop of a much higher density of other subtypes. Alternatively, loss of sensory input may have different effects on PV+ neurons in the two species.

Our finding that PV+ neuron soma area is reduced by ∼25 % at 7-14 days post-MMZ but recovers to baseline by 35 days, matching the time courses of OSN repopulation in the OE and of changes in TH+ neuron density, is intriguing. Indeed, naris occlusion in rats also significantly reduced PV+ soma area^4^. There is further precedent for activity-dependent changes in soma area in the olfactory, visual and auditory systems across species. In adult rats, mitral cell soma area decreased after 10 weeks of either deprivation of normal colony odor via rapid deodorized air flow, or constant exposure to cyclohexanone^71^. PV+ relay neurons in the cynomolgus monkey lateral geniculate nucleus also have reduced soma area after chronic (14 month) loss of optic nerve input in a glaucoma model^72^. Finally, soma area of neurons in auditory brainstem nuclei decreases within 24 hours of cochlear ablation or tetrodotoxin (TTX)-mediated silencing of the vestibulocochlear nerve in gerbils^73^. Of particular interest, soma area returned to baseline when activity was restored for 7 days following TTX-mediated silencing in this model, matching the recovery of OB PV+ soma area that we saw. However, what none of these studies addressed was the functional consequences of reduced soma area.

Input capacitance is directly proportional to membrane surface area, while input resistance is inversely proportional to membrane surface area^74^. Hence, the 33 % decrease in soma area that we found for PV+ neurons at 7 days post-MMZ would result in reduced capacitance and increased input resistance, which would be expected to increase excitability. This was corroborated by our integrate-and-fire model, which suggested a reduced rheobase and increased firing rate for PV+ neurons following MMZ treatment. This could markedly increase inhibition of OB output neurons^32^. However, while sensory input is absent or weak during the early stages of recovery, as after MMZ treatment, this would predominantly act to dampen spontaneous activity of OB output neurons. In addition, an important caveat is that we did not determine whether MMZ treatment also alters the dendritic morphology of PV+ neurons, which could also impact their excitability. Directly testing the impact of 7-14 days of MMZ treatment on both PV+ neuron excitability and PV-mediated inhibition of OB output neurons is an interesting future direction.

### Temporal pattern of changes in CR+ neuron density following MMZ treatment

Our most striking finding in this study is the transient and laminar-specific increase in GL and EPL CR+ neuron density at 7 days post-MMZ, which is no longer seen at 14 days post-MMZ. There are several potential mechanisms that could account for this transient increase. The first possibility is enhanced integration of CR+ neurons in the GL and EPL. While 7 days is very short for new SVZ-derived neurons to reach the GL, with 90 % of newborn juxtaglomerular neurons retrovirally labeled in the RMS arriving after 7 days post-injection^64^, CR+ neurons are also generated in the OB core^36,37^, enabling them to reach the GL within 7 days. Indeed, we found a 2-fold increase in the density of newborn EdU+CR+ neurons in the GL and EPL at 7 days post-MMZ, suggesting that OB core neurogenesis may be upregulated. However, the increase in the absolute number of CR+EdU+ neurons was very small (6.7 mm^-2^ in the GL) relative to the overall increase in CR+ neuron density (861 mm^-2^ in the GL). A single EdU injection one day after MMZ treatment may label only a fraction of the additional CR+ neurons that are being added, and EdU-mediated toxicity^75^ could also reduce the fraction of the newly added population that is labeled. However, it seems unlikely that we labeled less than 0.8 % of the newly generated CR+ neurons that reached the GL by 7 days post-MMZ. Furthermore, OB core neurogenesis generates similar proportions of CR+ PGCs and GCs^36,37^, but GCL CR+ neuron density was not increased at 7 days post-MMZ. Hence, we concluded that enhanced neurogenesis could not account for the rapid increase in CR+ density in the GL and EPL following MMZ treatment.

A second possibility is that JGN fate specification was altered. We found no difference in the proportion of EdU-labeled cells that expressed CR, suggesting that fate determination of neurons generated immediately after MMZ treatment is not. A more systematic study of newborn neurons pre-labeled with EdU at different time points prior to MMZ treatment would be required to conclusively answer the question of whether neuroblasts that were already en route to their final destination preferentially differentiated into CR+ PGCs. However, it is notable that a previous study found no difference in the proportions of CR+, CB+ and TH+ cells amongst neurons labeled with BrdU either before or after naris occlusion^5^, suggesting that fate determination is unlikely to be altered by loss of sensory input.

Finally, we considered whether CR expression might increase, such that cells that were previously undetectable by immunohistochemistry now expressed detectable levels of CR. Indeed, mean CR fluorescence intensity increased significantly in the GL and EPL. Hence, we concluded that increased CR expression in GL and EPL neurons is a significant contributor to the transiently increased CR+ neuron density observed 7 days post-MMZ. What are the functional effects of increased CR expression? As a hexa-EF-hand calcium binding protein with high cooperativity, CR buffers calcium, especially at high intracellular calcium concentrations, and may also function as a calcium sensor, undergoing calcium-dependent conformational changes^76^. CR knockout increased presynaptic release probability in cerebellar parallel fibers^77^ and increased excitability in cerebellar granule cells^78^. However, the effects of increased CR expression on CR+ PGCs, which are poorly connected with the local network and rarely fire action potentials^46,47^, are difficult to predict. Interestingly, it has been proposed that CR+ PGCs remain in an immature state, perhaps providing a reserve population that can be recruited when required by the circuit^47^. Hence, it is intriguing to speculate that increased CR expression may be indicative of greater integration of CR+ PGCs into the glomerular network in responses to the loss of sensory input induced by MMZ treatment.

At later time points post-MMZ, CR+ neuron density in the GL, EPL and GCL showed the same pattern of changes, with a decrease between 7 and 14 days and subsequent rebound at 35 days. The decrease between 7 and 14 days may reflect reduced CR expression or low-level cell death of CR+ neurons; distinguishing between these possibilities would require the use of CR-Cre mice. The increase in CR+ density between 14 and 35 days is mostly likely due to incorporation of newly generated CR+ neurons, as we also observed a positive trend in CR+ neuron density in saline-treated mice over the time course of our experiment (i.e. between 9 and 13 weeks of age). This suggests that CR+ neurogenesis may be additive over the lifespan of a mouse, as has been described for DA neurons^79^.

## Conclusion

This is the first study to assess changes in OB interneuron density following MMZ-mediated OSN ablation of sensory input. We found cell type- and laminar-specific changes in OB interneuron populations. Most notably, the soma area of PV+ neurons was transiently reduced at 7-14 but not 35 days post-MMZ, and the density of CR+ neurons in the GL and EPL, but not the GCL, was transiently increased only at 7 days post-MMZ. This highlights the range of activity-dependent plasticity mechanisms employed within OB circuits. It also raises important questions about the roles played by two very understudied types of neurons, CR+ PGCs and PV+ EPL interneurons, in the regenerating OB.

